# Empowering rare variant burden-based gene-trait association studies via optimized computational predictor choice

**DOI:** 10.1101/2021.09.20.459182

**Authors:** Da Kuang, Roujia Li, Yingzhou Wu, Jochen Weile, Robert A. Hegele, Frederick P. Roth

## Abstract

**Background:** Causal gene/trait relationships can be identified via observation of an excess (or reduced) burden of rare variation in a given gene within humans who have that trait. Although computational predictors can improve the power of such ‘burden’ tests, it is unclear which are optimal for this task.

**Method:** Using 140 gene-trait combinations with a reported rare-variant burden association, we evaluated the ability of 20 computational predictors to predict human traits. We used the best-performing predictors to increase the power of genome-wide rare variant burden scans based on ∼450K UK Biobank participants.

**Results:** Two predictors—VARITY and REVEL—outperformed all others in predicting human traits in the UK Biobank from missense variation. Genome-scale burden scans using the two best-performing predictors identified 1,038 gene-trait associations (FDR < 5%), including 567 (55%) that had not been previously reported. We explore 54 cardiovascular gene-trait associations (including 15 not reported in other burden scans) in greater depth.

**Conclusions:** Rigorous selection of computational missense variant effect predictors can improve the power of rare-variant burden scans for human gene-trait associations, yielding many new associations with potential value in informing mechanistic understanding and therapeutic development. The strategy we describe here is generalizable to future computational variant effect predictors, traits and organisms.

## Background

DNA sequencing in research and clinical genetic diagnostics have discovered millions of human genetic variants. Of particular interest are coding variants, which can alter protein functions and thus contribute to human diseases [1]. Many common coding variants have been causally linked to human diseases through genome-wide association studies (GWAS) [2]. Making causal associations between human traits and specific rare variants (which account for the vast majority of human genetic variants) is more difficult, given the limited statistical power of association methods and current cohort sizes [3]. However, by aggregating many variants into a single ‘pseudo-variant’ [4] that measures the overall burden of target gene variation within a subset of people, the burden of variation within people with a trait (cases) and those without a trait (controls) can be measured and compared via a ‘burden test’ [5]. Human genes can thus be confidently associated with traits, despite uncertainty about the pathogenicity of any specific aggregated variant [6, 7].

Burden tests can aggregate all observed coding variants, but neutral variation that is equally prevalent in both cases and controls can obscure associations between traits and the burden of gene-damaging variation. Alternatively, burden tests can consider only “qualifying” variants that are more likely to be pathogenic.

Qualification could be based on clinical variant interpretation frameworks [8, 9] that provide high-quality annotations of variant pathogenicity, but these have been applied to less than 2% of human coding variants observed thus far [10]. By contrast, computational variant effect predictors offer a readily available means of estimating variant pathogenicity. Widely used computational variant effect predictors developed over the last two decades include PROVEAN [11], PolyPhen-2 [12] and SIFT [13]. More recent variant effect predictors, e.g. EVE [14], have benefited from advances in deep learning. Some ‘meta-predictors’, e.g. REVEL [15] and VARITY [16], have achieved great performance in large part by combining the results of many evidence sources, including the results of other prediction algorithms.

The use of computational predictors to select qualifying variants has become common in burden testing [17–20]. The performance of computational variant effect predictors has the potential to affect the power and accuracy of burden testing, but it remains unclear how to choose the best predictor(s) for this purpose. Although several studies have benchmarked the performance of computational predictors [21–23], reliable assessment has many inherent challenges. Chief among these is the risk of circularity. Where “ground truth” test data have previously been used in training, performance estimates for a computational model may be artificially inflated [24].

To avoid circularity, Livesey and Marsh [21] assessed computational predictors using functional assay data. Well-established functional assays can provide strong evidence for or against pathogenicity [8, 25]. However, variant predictions could only be assessed for a few dozen proteins for which experimental “variant effect maps” were available. They also had limited ability to carry out unbiased evaluation for predictors (e.g., DeepSequence) that had used variant effect map data in model training. Although Livesey and Marsh found that variant effect maps were typically more accurate than predictors of pathogenicity, a variant effect map is currently available for only ∼1% of human disease-associated proteins [26].

An alternative source of evidence about variant effects, that has not previously been used in training predictors, is population-based cohorts of genotyped and phenotyped participants. For example, the UK Biobank [27] has assembled in-depth genetic and trait information for a prospective cohort of ∼500K participants. Whole-exome sequences for >450K participants have been released widely to researchers, and >7K data fields describing many human phenotypes are available for a large fraction of participants. Because variant effect predictors have not generally trained on these data, using the UK Biobank dataset as a source of “ground truth” human trait data sidesteps the risk of performance inflation due to circularity.

Here, we assessed computational predictors by examining the correspondence between variant scores and human phenotypes for 20 computational predictors, focusing on 140 gene-trait combinations previously reported in genome-wide rare variant burden studies [28–31], and identified VARITY and REVEL as best performing. A genome-wide rare variant burden scan with ∼450K participants in the UK Biobank cohort, using VARITY and REVEL to identify qualifying variants, revealed many more significant gene-trait associations than had previously been reported, and the scans using VARITY and REVEL more than doubled the number of reported burden-test-associated human gene-trait combinations. Although all scan results are provided, we further explore newly revealed burden-test associations involving cardiovascular traits.

## Results

### Assessing the performance of computational variant effect predictors

To examine the correspondence of computational variant effect predictors to human traits in UK Biobank participants, we: 1) identified 140 gene-trait combinations with reported burden-test associations in UK Biobank; 2) extracted rare missense variants from the corresponding genes; 3) obtained variant effect scores for these missense variants from 20 computational predictors; and 4) evaluated correspondence between predictor scores and human phenotypes. Figure 1 provides a schematic overview of the benchmarking framework.

**Figure 1.**
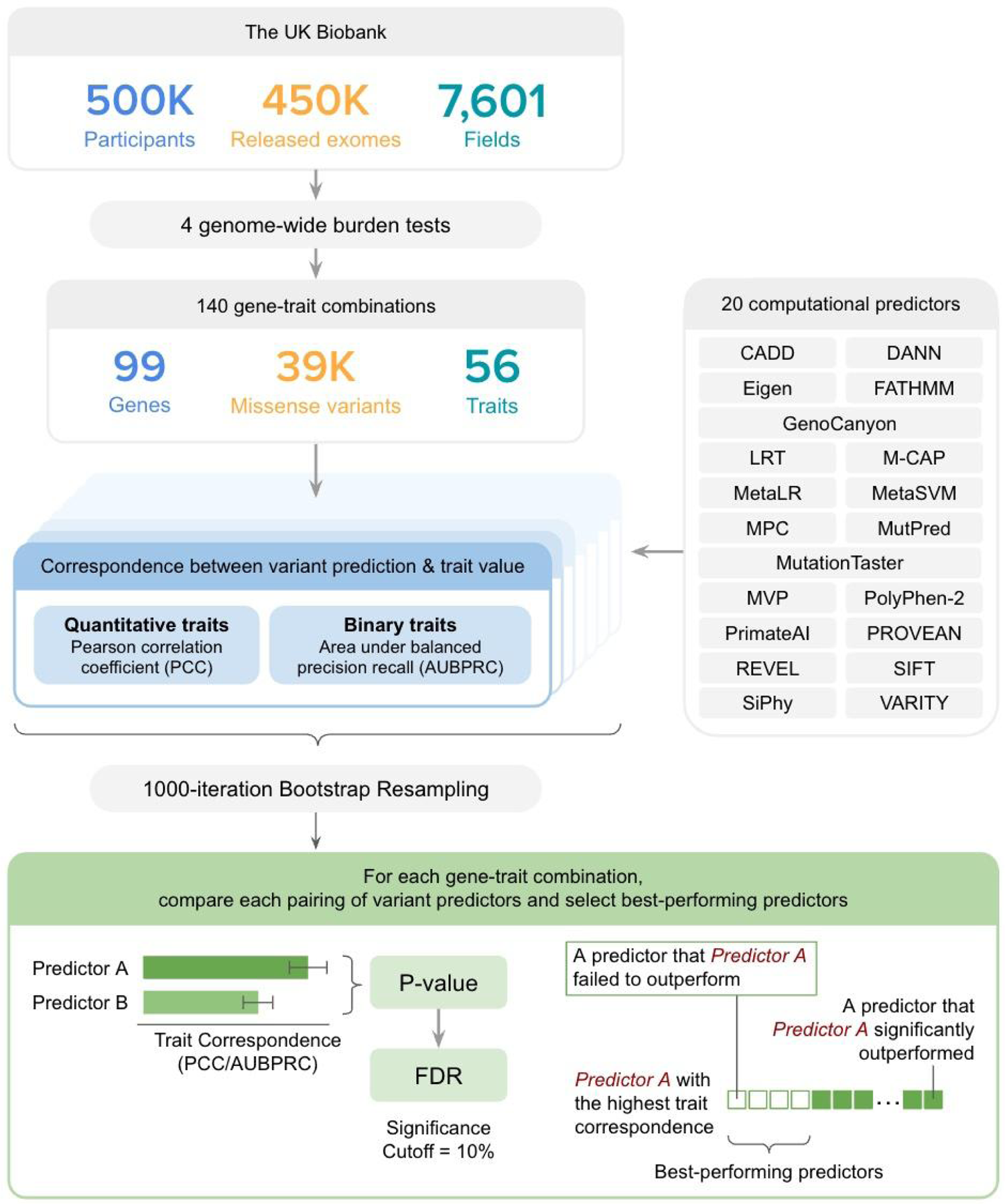
A schematic overview of the variant effect predictor benchmark. 140 gene-trait combinations were selected from the UK Biobank and used to assess the accuracy of 20 computational variant effect predictors (see Methods).

Based on four systematic burden testing studies performed using data from the UK Biobank cohort [28–31], we compiled a set of 140 gene-trait combinations. Table S1 lists the genes and the associated traits in the format of the UK Biobank field ID (FID). Of the 99 trait-associated genes, 73 (74%) were associated with only one trait. The remaining 26 genes were linked to multiple traits. For example, *LDLR*, a gene that encodes the low-density lipoprotein (LDL) receptor and is causal for autosomal dominant familial hypercholesterolemia (FH [MIM: 143890]) [32], was previously found to be associated with five traits: 1) blood LDL cholesterol level (mmol/L; FID: 30780); 2) self-reported high cholesterol (FID: 20002-1473); 3) taking atorvastatin (a cholesterol-lowering drug; FID: 20003-1141146234); 4) taking any cholesterol-lowering medication (FID: 6153-1); and 5) atherosclerotic heart disease of native coronary artery (FID: 41270-I25.1).

Having obtained whole-exome sequencing data released for >450K participants in the UK Biobank cohort (transfer of human data was approved and overseen by The UK Biobank Ethics Advisory Committee [Project ID: 51135]), we extracted coding single-nucleotide variants (SNVs) for the 99 associated genes. We focused all subsequent analyses on the subset of rare variants with a minor allele frequency (MAF) < 0.1%. Because the clinical annotation of “variant of uncertain significance” (VUS) is applied especially often to rare variants [33], we focused on predictor performance for rare variants. To determine whether the variant alters the amino acid sequence of the encoding protein, we mapped each variant to the canonical transcript of its corresponding gene in the Ensembl and CCDS database [34, 35], yielding an evaluation set of 39,696 rare human missense variants from 428,517 UK Biobank participants.

For each rare human missense variant, we obtained variant effect scores from 20 computational variant effect predictors (see Table S2 for a complete list of predictors compared in this study). Some predictors (e.g. PROVEAN) assign low scores to predicted-damaging variants, while others (e.g. PolyPhen-2) assign high scores to such variants. To better compare rankings from different predictors, we negated scores of the former type (i.e. low scores for predicted-damaging variants) so that the highest scores always corresponded to the variants that were predicted most strongly to be damaging.

We assessed, for each predictor and each gene, whether the predictor offered enough predictions for that gene to warrant inclusion in the comparison. Of the 20 predictors we examined, only 13 (65%) provided predictions for every missense variant in our evaluation set. However, for >90% of the genes of interest, all predictors provided scores for at least ten variants. Table S3 lists genes in which a predictor made predictions for less than ten variants. To ensure sufficient evidence to evaluate performance, predictors that made <10 predictions in a given gene were not included in the predictor performance assessment for that gene. To reduce the effect of extreme values on performance evaluation, we applied a floor and ceiling at the 5^th^ and 95^th^ percentile predictor scores, respectively, and applied a transformation giving all variant effect predictors the same 0-1 range of score values.

Next, we deployed two approaches to assess the performance of variant effect predictors, depending on whether the trait was binary or quantitative.

For gene-trait combinations with binary traits including categorical traits that could be simplified and considered binary, we applied an area under the balanced precision-recall curve (AUBPRC) approach. Because binary trait measurements were made on the participant level, i.e. one measurement per participant in the UK Biobank cohort, we sought a participant-level summary of predicted variant effects. Therefore, we summed the total predictor score for all missense variants observed in each gene of interest in each participant. We note that this approach essentially models variant effects as being additive, e.g., two mildly damaging variants (score = 0.5) combined will show a more damaging effect (total score = 1.0). While other approaches exist to model multiple variants, given that, on average, only ∼1% of participants had more than one variant in each gene (Figure S1), more sophisticated models would be unlikely to substantially affect our results.

Illustrating our evaluation of the extent to which computational variant effect scores can predict phenotypic traits, Figure 2 shows results for UK Biobank participants who carry rare missense variants in the *LDLR* gene. Participants with missense variants that have a predicted-damaging total missense variant impact (participant-centric score ≥ 0.9, using VARITY as the source of variant scores) are three times as likely to have been prescribed cholesterol-lowering medication. This result aligns with the finding that participants with clinically pathogenic *LDLR* missense variants have an elevated risk for hypercholesterolemia [36] and, as a result, are more likely to receive cholesterol-lowering medication, e.g. a statin or proprotein convertase subtilisin kexin type 9 (PCSK9) inhibitor.

**Figure 2.**
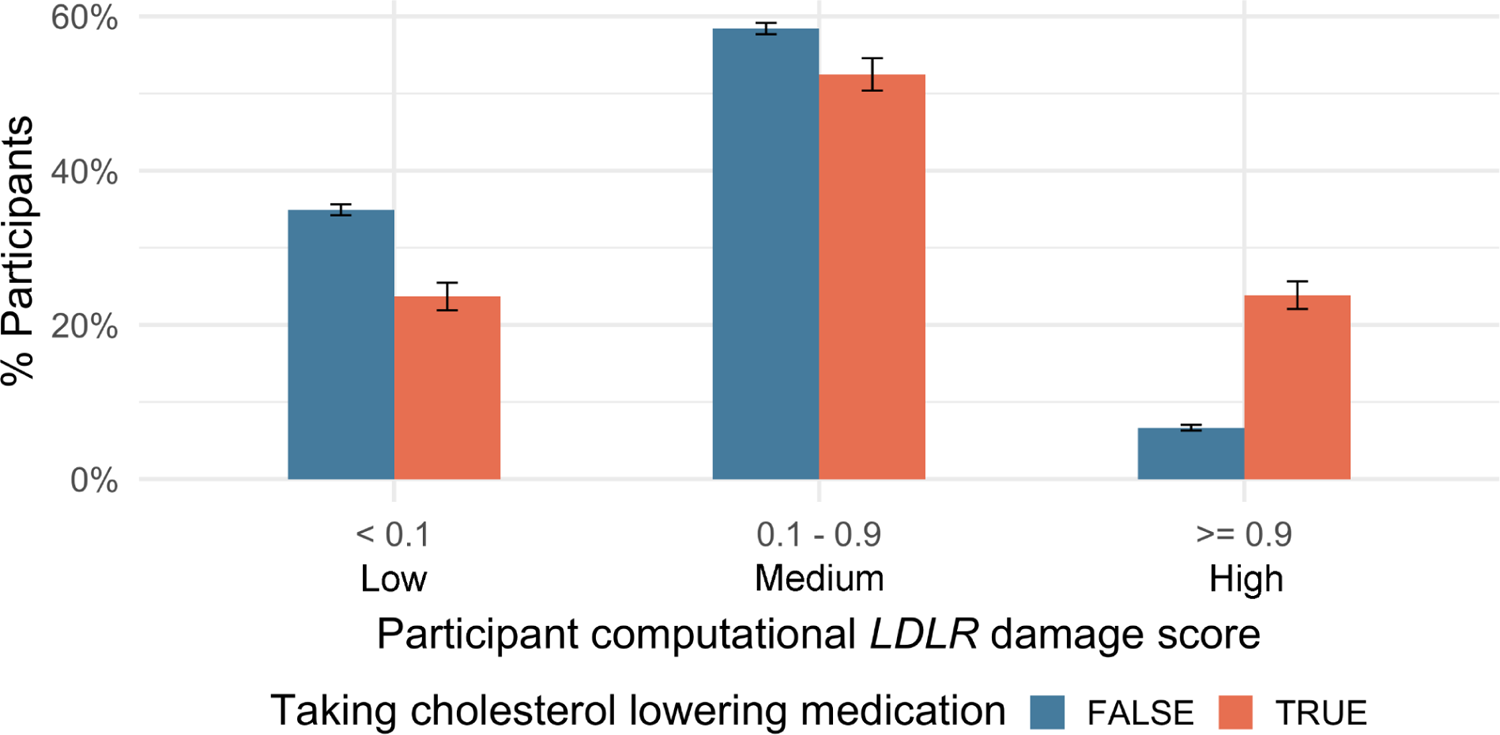
Percentages of UK Biobank participants with different participant computational *LDLR* damage scores for participants. This example examined participants either taking or not taking cholesterol-lowering medications, as well as rare missense variants in *LDLR*. Computational *LDLR* damage scores were used in which higher scores indicate more damaging variants. Error bars represent standard errors.

For every gene-trait combination and each predictor, we analyzed the tradeoff between precision (i.e. the fraction of participants above a given total variant impact score threshold that had the trait value associated with heightened rare-variant burden) and recall (i.e. the fraction of all participants with the rare-variant-burden-associated trait value that were detected at a given total variant impact score threshold). More specifically, because precision depends on the prior probability of the trait value, we evaluated balanced precision, which represents precision where the prior probability of the trait value is 50% [16]. This enabled us to evaluate, for each combination of gene and binary trait, the AUBPRC for each computational predictor. Variant predictors were considered better performing if they had a significantly higher AUBPRC.

Because AUBPRC analysis is only appropriate for binary traits, for quantitative traits we used the Pearson Correlation Coefficient (PCC) to assess the correspondence between variant impact score and trait value. Where multiple participants carried the same variant, we averaged their quantitative trait values. Variant predictors were considered better performing if they had a significantly higher PCC.

Thus, for each gene-trait combination, we obtained either a PCC or AUBPRC measure of performance. To estimate uncertainty in each performance measure, we carried out 1000-iteration bootstrap resampling in which variants of a given gene were resampled with replacement, and PCC or AUBPRC values were re-calculated for each sample. For each gene-trait combination, this yielded a distribution of either PCC or AUBPRC values for each computational predictor. From each of these distributions, we extracted a mean and a 95% confidence interval (CI) that reflects our uncertainty in the performance measure.

To illustrate this approach, Figure 3 shows AUBPRC and PCC values and CIs for each of 20 computational predictors for one binary (top panel) and one quantitative (bottom panel) gene-trait combination involving *LDLR*. Here, the numerically top-performing variant effect predictor was VARITY. To assess whether numerical differences were statistically significant, we computed empirical p-values between every pair of computational predictors for each gene-trait combination. Then, to correct for having tested multiple hypotheses (one for each pairwise prediction comparison), we used the distribution of p-values for each gene-trait combination to derive corresponding FDR values. We considered a predictor Y to significantly outperform a predictor X if the comparison yielded an FDR < 10%. FDRs for all predictor pairs are shown for the gene-trait combination of *LDLR* with blood LDL level (mmol/L) (Figure 4). Although VARITY exhibited the highest PCC, three predictors (VARITY, MutPred and REVEL) were statistically indistinguishable, so that these three were each considered to be a “top” predictor for this gene-trait combination.

**Figure 3.**
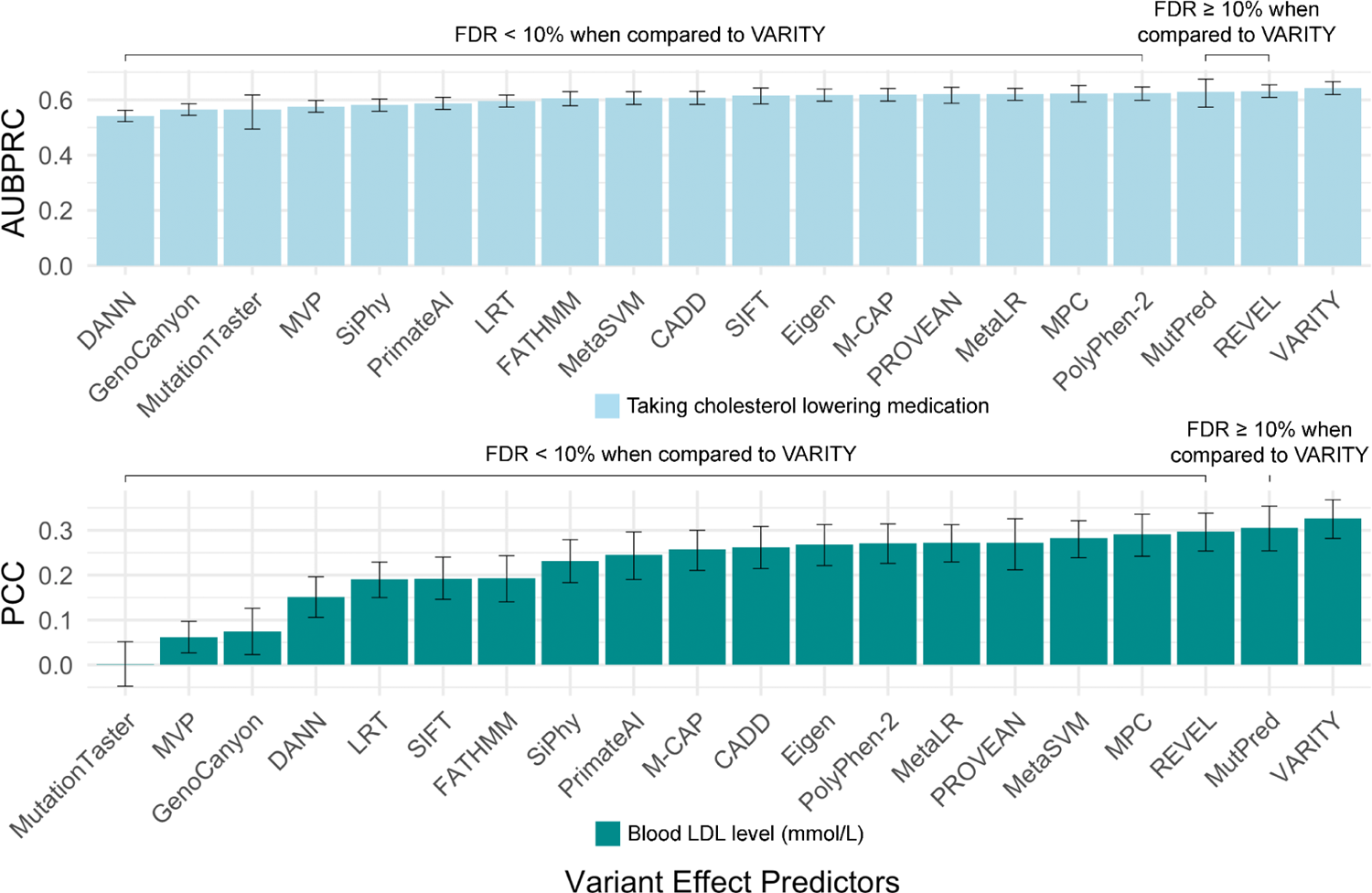
The performance of 20 computational methods in predicting two cholesterol-related phenotypes based on presence of rare *LDLR* missense variants. Performance comparisons used mean AUBPRC for binary trait “taking cholesterol-lowering medication” (top panel) and mean PCC for quantitative trait “blood LDL level (mmol/L)” (bottom panel), respectively. In both panels, significance indicators show whether predictor performance was significantly lower (FDR < 10%) than VARITY, the numerically best performing predictor. Error bars indicate 95% confidence intervals.

**Figure 4.**
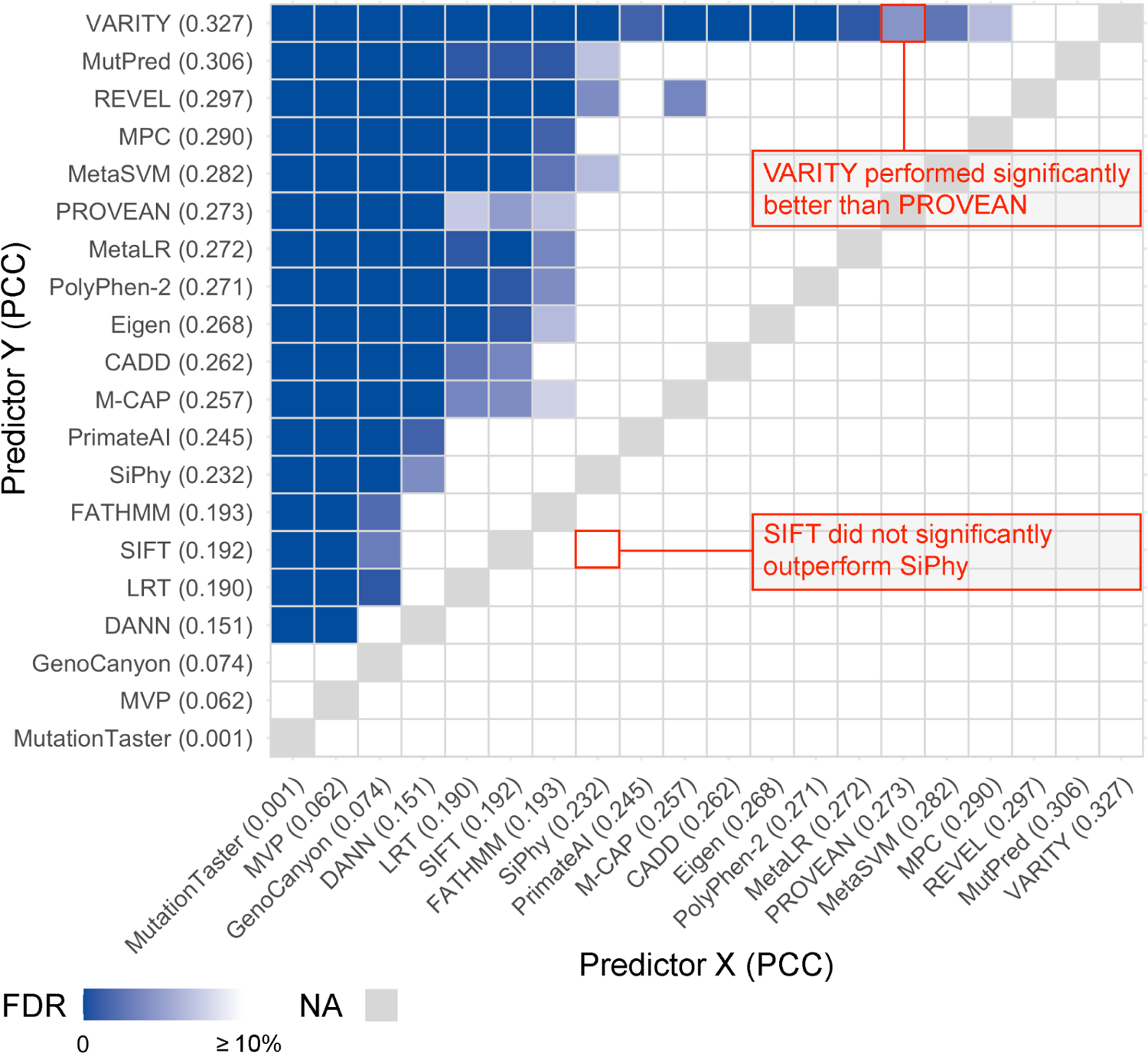
Example summary of pairwise comparisons between computational predictors, evaluating how well predictor scores for rare *LDLR* missense variants correspond to participant’s blood LDL level (mmol/L). Variant effect predictors are ranked top-to-bottom and right-to-left based on decreasing performance (PCC). Comparisons in which one predictor Y outperforms another predictor X (with FDR < 10%) are indicated in blue.

To summarize similar comparisons over all 140 gene-trait combinations, we counted the number of combinations in which a computational predictor was considered best-performing (Figure 5). These results showed that VARITY performed best across the largest number of gene-trait combinations: 135 (96%) of 140, while the next best predictor, REVEL, was a top predictor for 131 (94%) of 140 gene-trait combinations. The performance among the remaining 18 predictors ranged from 60% to 85%, with a mean of 76%. The best variant predictor, according to this ranking, was VARITY [16], followed by REVEL [15] and Eigen [37]. Because VARITY and REVEL were statistically indistinguishable (FDR = 22%; Wilcoxon signed-rank test), we considered both VARITY and REVEL to be the best performing in this evaluation.

**Figure 5.**
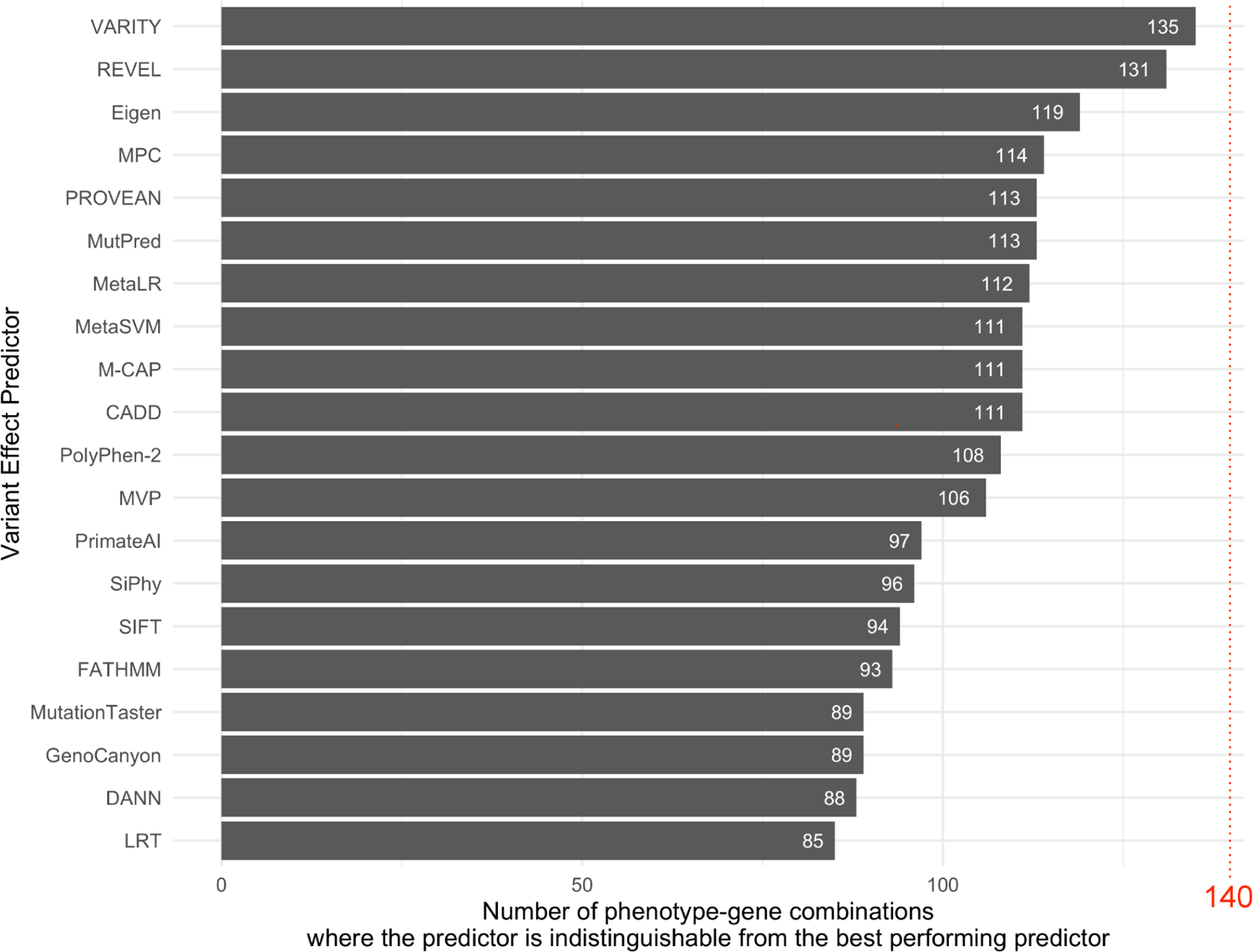
The number of gene-trait combinations for which each predictor is either best-performing or indistinguishable from best-performing. The red line highlights the maximum value based on the 140 gene-trait combinations considered in this study.

Despite excellent reported performance [14], the EVE predictor provided scores for many fewer variants than other methods, and we could evaluate it for only 71 (51%) of the 140 gene-trait combinations considered here. We therefore also established a smaller benchmark set of 57 gene-trait combinations for which all predictors (including EVE) could be evaluated (Figure S2A). In this evaluation, VARITY was a top predictor for 50 (88%) out of 57 gene-trait combinations, while EVE and REVEL were top predictors for 48 (84%). Despite being numerically better in this evaluation, VARITY’s performance was statistically indistinguishable from EVE and REVEL (Figure S2B), while VARITY, REVEL and EVE each significantly outperformed all other methods evaluated. Given that EVE provided fewer variant scores, we opted to use VARITY and REVEL in all subsequent analyses.

### Performing a genome-scale rare variant burden scan using top predictors

Hypothesizing that top-performing computational predictors could serve to increase the power of burden tests, we separately employed VARITY and REVEL for genome-wide burden scans based on rare (MAF < 0.1%) variants in 428K UK Biobank participants of European descent (see Figure 6 for an overview of burden scans). We selected 14,107 binary and quantitative traits (Table S4) available for the UK Biobank cohort, casting a wide net for disease-related traits. For each given trait, participants with the trait were designated as a “case” and those without as a “control”. Quantitative traits were first converted to binary traits using trait-specific thresholds (see Methods for details).

**Figure 6.**
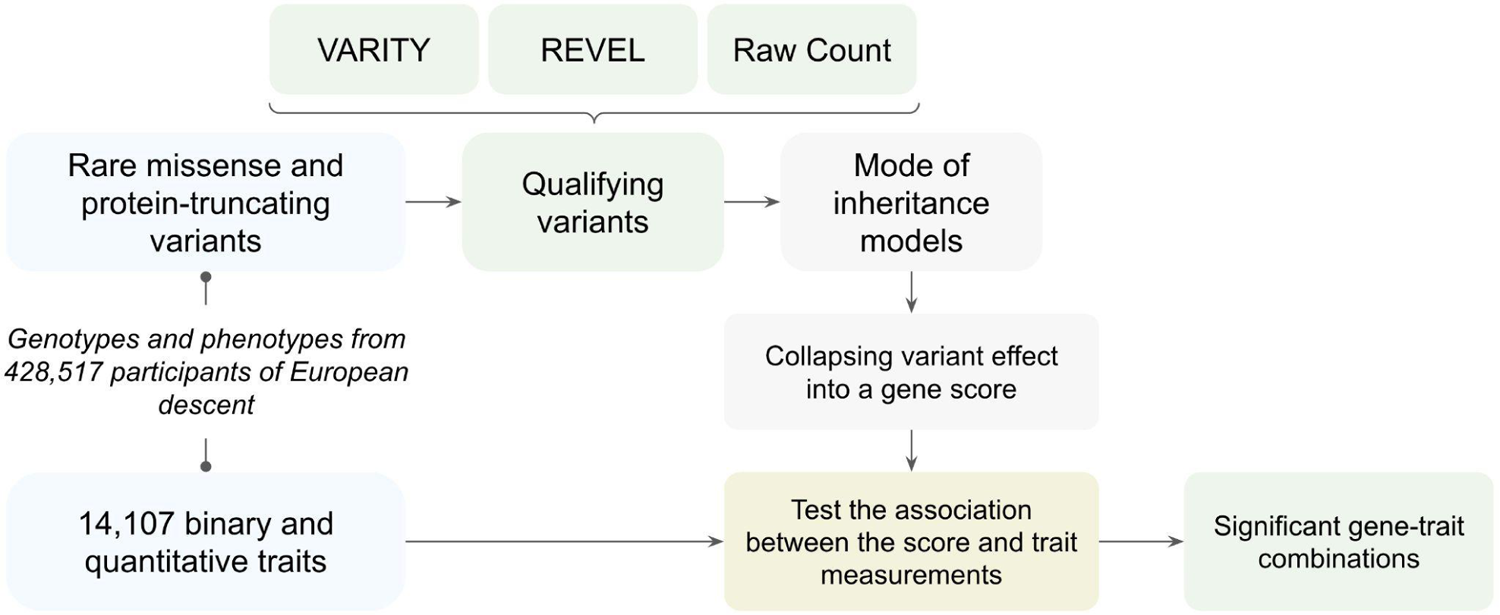
A schematic overview of the burden test. This burden test was based on 14,107 traits and the rare missense and protein-truncating variants from 428K participants in the UK Biobank cohort (see Methods).

Next, using both VARITY and REVEL, we identified qualifying rare missense variants from UK Biobank (see Methods). On average, 5% of unique rare missense variants (and 4% of all instances of rare missense variants) observed in the UK Biobank were considered to be qualifying variants. In addition, all protein-truncating rare variants were included as qualifying variants. We also included a “No Predictor” control, considering all rare missense variants as qualifying variants.

Because human diseases have modes of inheritance that range from dominant to recessive, our method modelled these two modes of inheritance separately. In the dominant model, participants were counted as carrying a variant in a gene if they had at least one qualifying missense variant in that gene, while the recessive model required the presence of at least two qualifying variants. Future applications of the recessive model might require at least one qualifying variant to be present in each allele, but this would require the availability of phased whole exomes.

To identify gene-trait combinations for which the gene exhibits a significantly elevated (or decreased) rare-variant burden in those with the trait, we first established a null distribution by randomly shuffling participant identifiers as previously described [38, 39]. For each gene-trait combination, a nominal p-value was calculated with Fisher’s exact test. To correct for multiple hypothesis testing, FDRs were calculated based on the above-described null distribution. Gene-trait combinations found to be significantly associated under each mode of inheritance model, together with the nature (risk vs. protective) and effect size (odds ratio) are provided in Table S5.

The burden scans using REVEL and VARITY identified 657 and 551 significant gene-trait combinations, respectively, while the burden scan without computational predictors (“No Predictor”) only identified 232 gene-trait combinations. A total of 1,038 significant gene-trait combinations were identified by at least one of the three methods used to determine qualifying variants, corresponding to 0.004% of the 26,984,770 gene-trait combinations considered. Of these 1,038 gene-trait combinations, 222, 295 and 170 were uniquely found by VARITY, REVEL and “No Predictor”, respectively. Thus, using computational predictors to call qualifying variants increased the sensitivity of detecting significant gene-trait combinations.

As an initial disease-focused exploration of the burden scan results, we examined the results for cardiovascular traits. Table 1 shows the 52 cardiovascular trait associations identified (FDR < 5%) by our burden scan. Of these, 38 (73%) had been reported in another burden test based on 450K UK Biobank participants [39], and an additional 5 (10%) had previous support from the literature. Indeed, our findings included many known associations, including some of high therapeutic and diagnostic value. For example, the scan correctly reported the well-known effect of *PCSK9* rare variants on lowering the blood LDL cholesterol level. This protective effect prompted the development of *PCSK9* inhibitors, a relatively recent therapeutic avenue for lowering the cholesterol level in patients [40]. Another example was the gene-trait combination of *APOC3* with blood triglyceride level. Rare variants in *APOC3* are known to lower blood triglyceride levels [41], and the APOC3 inhibitor volanesorsen recently completed a phase 3 clinical trial with promising results both in reducing the blood triglyceride level and possibly the risk of cardiovascular outcomes [42]. Other examples of genotype-phenotype correlations that are supported by mendelian genetics include the associations between HDL cholesterol and *ABCA1*, *LCAT*, *APOA1, SCARB1* and *CETP*, and between triglycerides and *LPL* and *APOA5* [43].

**Table 1.** 52 cardiovascular gene-trait combinations we found to have significant rare variant burden. A checkmark (✓) indicates that a gene-trait combination was found to be significant by a certain method (VARITY/REVEL/No Predictor) calling qualifying variants (see Methods for detail). The literature support column lists published evidence (if any) supporting each of the gene-trait combinations. Also listed is whether a given gene-trait combination was novel relative to Wang et al., a state-of-the-art burden test with 450K UK Biobank participants [39]. If a gene-trait combination has closely related significant combinations in Wang et al., it’s marked as “=”, otherwise, it is marked as “✓”.

| Gene | Trait | Qualifying-variant method yielding significant burden |  |  | Literature support | Novel relative to Wang et al. (burden test with 450K UK Biobank data) |
| --- | --- | --- | --- | --- | --- | --- |
|  |  | VARITY | REVEL | No Predictor |  |  |
| *APOA5* | Triglycerides (mmol/L) | ✓ | ✓ |  | *APOA5* variants modulate levels of triglycerides [72] | = |
|  | NMR Total Triglycerides (mmol/L) | ✓ | ✓ |  | *APOA5* variants modulate levels of triglycerides [72] | = |
| *APOB* | LDL cholesterol (mmol/L) | ✓ | ✓ |  | ApoB is a component of LDL [73] | = |
|  | NMR LDL cholesterol (mmol/L) | ✓ | ✓ |  | ApoB is a component of LDL [73] | = |
|  | Cholesterol (mmol/L) | ✓ | ✓ |  | ApoB forms lipoproteins which carry cholesterol [74] | = |
|  | NMR Total Cholesterol (mmol/L) | ✓ | ✓ |  | ApoB forms lipoproteins which carry cholesterol [74] | = |
|  | NMR Total Cholesterol minus HDL Cholesterol (mmol/L) | ✓ | ✓ |  | ApoB forms lipoproteins which carry cholesterol [74] | = |
|  | Apolipoprotein B (g/L) | ✓ | ✓ |  | *APOB* encodes ApoB | = |
|  | NMR Apolipoprotein B (g/L) | ✓ | ✓ |  | *APOB* encodes ApoB | = |
|  | Triglycerides (mmol/L) | ✓ | ✓ |  | ApoB forms triglyceride-rich lipoproteins [75] | = |
|  | NMR Total Triglycerides (mmol/L) | ✓ | ✓ |  | ApoB forms triglyceride-rich lipoproteins [75] | = |
|  | HDL Cholesterol (mmol/L) | ✓ |  |  | ApoB forms lipoproteins which carry cholesterol [74] | = |
| *ABCA1* | Apolipoprotein A (g/L) | ✓ | ✓ |  | ABCA1 mediates the formation of ApoA [76] | = |
|  | NMR Apolipoprotein A1 (g/L) | ✓ | ✓ |  | ABCA1 mediates the formation of ApoA [76] | = |
|  | NMR HDL cholesterol (mmol/L) |  | ✓ |  | ABCA1 mediates the efflux of cholesterol [76] | = |
|  | HDL cholesterol (mmol/L) |  | ✓ |  | ABCA1 mediates the efflux of cholesterol [76] | = |
| *ZNF226* | LDL cholesterol (mmol/L) | ✓ |  |  |  | ✓ |
| *LCAT* | Apolipoprotein A (g/L) | ✓ |  |  | LCAT enables maturation of HDL (formed by ApoA) [77] | = |
|  | HDL cholesterol (mmol/L) | ✓ |  |  | LCAT enables maturation of HDL [77] | = |
|  | NMR HDL cholesterol (mmol/L) |  | ✓ |  | LCAT enables maturation of HDL [77] | = |
| *PCSK9* | NMR Apolipoprotein B (g/L) | ✓ | ✓ |  | PCSK9 prevents ApoB from degradation [78] | = |
|  | LDL cholesterol (mmol/L) | ✓ | ✓ |  | PCSK9 inhibitors lower blood LDL [40] | = |
|  | NMR Total Cholesterol minus HDL Cholesterol (mmol/L) | ✓ | ✓ |  | PCSK9 inhibitors lower blood LDL [40] | = |
|  | NMR Total Cholesterol (mmol/L) | ✓ | ✓ |  | PCSK9 inhibitors lower blood LDL [40] | = |
|  | Cholesterol (mmol/L) |  | ✓ |  | PCSK9 inhibitors lower blood LDL [40] | = |
| *LDLR* | NMR LDL cholesterol (mmol/L) | ✓ | ✓ |  | LDLR removes LDL from the blood [79] | = |
|  | Treatment: atorvastatin | ✓ |  |  | Atorvastatin increases LDLR expression [80] | ✓ |
|  | LDL cholesterol (mmol/L) | ✓ |  |  | LDLR removes LDL from the blood [79] | = |
|  | Cholesterol (mmol/L) | ✓ |  |  | LDLR removes cholesterol from the blood [79] | = |
|  | NMR Total Cholesterol (mmol/L) |  | ✓ |  | LDLR removes cholesterol from the blood [79] | = |
|  | NMR Total Cholesterol minus HDL Cholesterol (mmol/L) |  | ✓ |  | LDLR removes cholesterol from the blood [79] | = |
|  | Apolipoprotein B (g/L) |  | ✓ |  | LDLR removes ApoB from the blood [79] | = |
| *CETP* | HDL cholesterol (mmol/L) | ✓ | ✓ |  | CETP inhibitors increase HDL level [81] | = |
|  | Apolipoprotein A (g/L) | ✓ | ✓ |  | Expression of *CETP* causes release of ApoA from HDL particle [82] | = |
| *PROC* | Self-reported illness: pulmonary embolism +/- deep venous thrombosis | ✓ |  |  | PROC controls blood clotting [83] | ✓ |
| *APOA1* | Apolipoprotein A (g/L) | ✓ | ✓ |  | *APOA1* encodes ApoA1 | = |
|  | HDL cholesterol (mmol/L) | ✓ | ✓ |  | ApoA1 forms HDL [84] | = |
| *SCARB1* | HDL cholesterol (mmol/L) |  | ✓ |  | *SCARB1* variants are linked to high HDL [85] | = |
| *AGPAT4* | Lipoprotein A (nmol/L) |  |  | ✓ | *AGPAT4* is involved in lipoprotein metabolism [86] | = |
| *PRSS12* | HDL cholesterol (mmol/L) |  | ✓ |  |  | ✓ |
| *PTEN* | HDL cholesterol (mmol/L) | ✓ | ✓ |  | Loss of *PTEN* increases blood cholesterol level [87] | ✓ |
| *GPR176* | Apolipoprotein B (g/L) | ✓ |  |  |  | ✓ |
| *TAAR8* | Self-reported illness: stroke |  |  | ✓ | 3-T1AM (*TAAR8* agonist) protects against brain damage upon stroke [88] | ✓ |
| *AGPAT4* | Lipoprotein A (g/L) |  |  | ✓ | *AGPAT4* is involved in lipogenesis and complex lipid synthesis [89] | = |
| *PGK1* | HDL cholesterol (mmol/L) | ✓ |  |  |  | ✓ |
| *HELZ* | Body mass index (BMI) | ✓ |  |  |  | ✓ |
| *CBL* | HDL cholesterol (mmol/L) | ✓ | ✓ |  |  | ✓ |
| *GAMT* | Lipoprotein A (g/L) |  | ✓ |  |  | ✓ |
| *DDX3Y* | HDL cholesterol (mmol/L) |  |  | ✓ |  | ✓ |
| *CT45A1* | HDL cholesterol (mmol/L) |  |  | ✓ |  | ✓ |
| *LPL* | Triglycerides (mmol/L) |  | ✓ |  | LPL defects are linked to higher triglycerides [90] | = |
| *KRT6C* | LDL cholesterol (mmol/L) |  |  | ✓ | [Indirect evidence] KRT6C interacts with KRT31 [91] which is GWAS-associated with LDL level [92] | ✓ |

The remaining 9 (17%) of the cardiovascular-related gene-trait combinations we found are novel, having neither been reported by previous burden scans nor described previously in the literature. One example is the protective effect found here for rare variants in *ZNF226* on LDL cholesterol levels. The zinc finger protein 226 (ZNF226), predicted to enable DNA-binding and transcriptional activation [44], has not previously been associated with coronary artery disease-related traits. Future studies might investigate the potential protective effect of rare variants in *ZNF226* and assess whether *ZNF226* can become a new therapeutic target for lowering cardiovascular risk factors.

Consistent with the overall analysis, the sensitivity of the burden scan for cardiovascular traits depended strongly on the variant effect predictor used. Although 24 (46%) of the 52 gene-trait combinations identified were found using both VARITY and REVEL, 11 (21%) were identified only with VARITY or REVEL, as compared with only 6 (11%) associations uniquely found by not using a predictor. Thus, consistent with the overall findings, VARITY and REVEL enabled improved sensitivity to detect cardiovascular trait associations.

Beyond cardiovascular traits, the VARITY- and REVEL-guided burden scans we performed found many disease associations that have not been previously described in other burden scans [39]. The 20 largest-effect disease associations that were novel relative to previously published rare-variant burden scans are shown in Table 2, together with literature citations (where available) that support these associations. Novel associations include the finding that *SLC25A5* is associated with “depression possibly related to childbirth” (FID: 20445), which is supported by a previous report linking altered *SLC25A5* expression to depression [45]. In another example, we found that *TET2* burden is associated with participants having received diagnostic puncture of bone (FID: 41200), which is expected because *TET2* is linked to clonal hematopoiesis disease [46], which can be diagnosed via bone marrow aspiration [47].

**Table 2.** Top 20 disease-related gene-trait associations that were novel relative to previously published rare-variant burden scans [39], ranked by the absolute log10 odds ratio. If a gene-trait combination was found to be significant by multiple methods (VARITY/REVEL/No Predictor), the absolute log10 odds ratios were first averaged prior to ranking. For genes associated with quantitative traits, the predicted effect of rare variant burden is provided. The literature support column lists published evidence (if any) supporting each of the gene-trait combinations.

| Gene | Trait | Effect size<br>(absolute log10<br>odds ratio) | Variant<br>effect on<br>trait | Literature support |
| --- | --- | --- | --- | --- |
| *SLC25A5* | Depression possibly related to childbirth | 1.66 |  | Altered *SLC25A5* expression is linked to depression [45] |
| *HBA1* | Mean corpuscular haemoglobin (pg) | 1.62 | Lowered | HBA1 deficiency is linked to anemia (OMIM: 613978) |
| *LRRTM4* | Neuroticism score | 1.38 | Increased | LRRTM4 regulates excitatory synapse development [93] |
| *UBXN6* | Ever had bowel cancer screening | 1.30 |  | UBXN6 is a potential predictor of postoperative recurrence in gastric cancer [94] |
| *ARPC1B* | Seen doctor (GP) for nerves, anxiety, tension or depression | 1.27 |  | ARPC1B might protect rats from chronic mild stress [95] |
| *THPO* | Platelet crit (%) | 1.22 | Lowered | Loss-of-function *THPO* variants are linked to increased platelet crit [96] |
| *TET2* | Operation: Diagnostic puncture of bone | 1.20 |  | TET2 is linked to hematopoietic disease [46] which is diagnosed via bone marrow aspiration [47] |
| *CDKN2A* | Diagnosis: Melanocytic naevi | 1.13 |  | *CDKN2A* variants are associated with melanocytic naevi [97] |
| *SMYD5* | Systolic blood pressure, manual reading (mmHg) | 1.11 | Lowered |  |
| *MAPRE1* | NMR Total Concentration of Branched-Chain Amino Acids (mmol/L) | 1.10 | Lowered |  |
| *HBA1* | Mean corpuscular volume (fL) | 1.10 | Lowered | HBA1 deficiency is linked to anemia (OMIM: 613978) |
| *JAK2* | Medication: vitamin c product | 1.04 |  | JAK2 function is induced by Vitamin C [98] |
| *BRCA2* | Operation: Total excision of breast | 1.04 |  | Prophylactic mastectomy can reduce breast cancer risk in people carrying *BRCA1/2* variants [99] |
| *GATA1* | Platelet distribution width (%) | 1.02 | Increased | GATA1 deficiency is associated with enlarged platelets [100] |
| *BORA* | Glycated haemoglobin (HbA1c) (mmol/mol) | 1.01 | Increased |  |
| *UBE2Z* | Platelet crit (%) | 1.00 | Lowered |  |
| *TRMT112* | NMR Phospholipids in Medium LDL (mmol/L) | 1.00 | Lowered |  |
|  | NMR Total Lipids in Medium LDL (mmol/L) | 1.00 |  |  |
| *CCR2* | NMR Triglycerides in HDL (mmol/L) | 1.00 | Lowered | CCR2 knockout mice have lower triglyceride levels [101] |
| *AL163195.3* | NMR Free Cholesterol in Very Large HDL (mmol/L) | 1.00 | Increased |  |

These burden scans also identified associations for many non-disease-related traits that had not been previously described in other burden scans. The 20 largest-effect non-disease trait associations that were novel relative to previously published rare-variant burden scans are shown in Table 3, together with literature citations supporting these associations, where available. We found several genes to have a burden of mutation associated with body mass related phenotypes, that are potential targets for therapeutic weight control. For example, *GSTK1* negatively correlates with the arm fat mass phenotype (FID: 23124), which is supported by literature evidence indicating that *GSTK1* is inversely correlated with obesity [48].

**Table 3.** Top 20 non-disease gene-trait associations that were novel relative to previously published rare-variant burden scans [39], ranked by the absolute log10 odds ratio. If a gene-trait combination was found to be significant by multiple methods (VARITY/REVEL/No Predictor), the absolute log10 odds ratios were first averaged prior to ranking. For genes associated with quantitative traits, the predicted effect of rare variant burden is provided. The literature support column lists published evidence (if any) supporting each of the gene-trait combinations.

| Gene | Trait | Effect size<br>(absolute log10<br>odds ratio) | Variant<br>effect on<br>trait | Literature support |
| --- | --- | --- | --- | --- |
| *SLC25A5* | Leg fat-free mass (right) (Kg) | 1.18 | Increased | SLC25A5 inhibits adipogenesis [102] |
|  | Arm predicted mass (left) (Kg) | 1.18 |  |  |
|  | Arm predicted mass (right) (Kg) | 1.16 |  |  |
| *PLAG1* | Arm fat-free mass (left) (Kg) | 1.16 | Lowered | PLAG1 knockout mice were ~30% smaller than wildtype mice [103]. |
| *SLC25A5* | Whole body water mass (Kg) | 1.15 | Increased | SLC25A5 inhibits adipogenesis [102] |
|  | Trunk predicted mass (Kg) | 1.15 |  |  |
|  | Trunk fat-free mass (Kg) | 1.15 |  |  |
| *MAPRE1* | NMR Total Concentration of Branched-Chain Amino Acids (mmol/L) | 1.10 | Lowered |  |
| *PLAG1* | Whole body water mass (Kg) | 1.10 | Lowered | PLAG1 knockout mice were ~30% smaller than wildtype mice [103]. |
|  | Whole body fat-free mass (Kg) | 1.08 |  |  |
| *EAPP* | Total sugars (g) | 1.02 | Lowered |  |
| *ZWINT* | NMR Docosaehxaenoic Acid (mmol/L) | 0.98 | Increased | ZWINT is linked to fatty acid degradation [104] |
| *MARVELD1* | Whole body water mass (Kg) | 0.95 | Increased |  |
| *SIGLEC6* | Age stopped smoking | 0.95 | Increased |  |
| *MARVELD1* | Whole body fat-free mass (Kg) | 0.95 | Increased |  |
| *GSTK1* | Arm fat mass (left) (Kg) | 0.94 | Lowered | GSTK1 level is inversely correlated with obesity [48] |
| *HSD3B7* | Trunk fat percentage | 0.91 | Increased |  |
| *CP* | Ever addicted to a behaviour or miscellaneous | 0.91 |  |  |
| *HSD3B7* | Body fat percentage | 0.91 | Increased |  |
| *KPNA1* | Born by caesarian section | 0.90 |  |  |

As illustrated by the exploration of cardiovascular traits, the increased-sensitivity burden scan results we provide here have the potential to nucleate many future explorations of disease mechanisms and therapeutic targets.

## Discussion

Although computational predictors of missense variant impact are known to increase the power of rare-variant burden tests for detecting gene-trait associations [5, 17], no previous study has systematically evaluated and compared the best predictors for this purpose. Therefore, for 20 computational predictors, we evaluated the strength of correspondence to human traits in UK Biobank participants, considering 140 gene-trait combinations for which a burden association had previously been shown.

Because none of the computational predictors we examined had been trained using UK Biobank data, our evaluation approach has the marked advantage of independence and avoidance of performance inflation that can arise when predictors are assessed using data on which they were previously trained. Although every predictor was best (or tied for best) for at least one gene-trait combination, counting the number of gene-trait combinations in which a predictor was best enabled an overall ranking.

It is interesting that the top three predictors overall – VARITY [16], REVEL [15], and Eigen [37] – are all meta-predictors, i.e., they combine prediction scores from other variant effect predictors. For meta-predictors, it can be challenging to establish “ground truth” sets of variants that had not been used for training any of the input predictors. That said, VARITY only exploited scores from other predictors that were unsupervised, i.e., made no direct use of variant pathogenicity annotations. The fourth-ranked predictor, MPC [49], exploited observations of depleted missense variation within particular sub-genic regions. That none of the top-three-ranked predictors leveraged this kind of information suggests the possibility that combining these approaches could yield still-better results.

One limitation of our study was that we did not evaluate all published predictors. This was in part due to the abundance of such predictors, but many predictors were excluded due to non-functional websites. However, we were able to examine variant effect predictors 1) that are widely used to predict variant effects (e.g. PolyPhen-2, PROVEAN) and 2) for which there is evidence of performance that exceeds other predictors (e.g. PrimateAI, REVEL, EVE). We suggest that this analysis should be repeated periodically to benchmark and test evaluate predictors as they emerge.

Another limitation of the predictor comparison study is that we did not consider correlations between traits. For example, *LDLR* was associated with multiple mutually-correlated traits, so our analysis was influenced by some genes and traits more than others. That said, body mass index (BMI; FID: 21001) was the most recurrent trait (appearing in 23 gene-trait combinations), and *LDLR* was the most recurrent gene (appearing in 5 gene-trait combinations). Thus, no one gene or trait dominated the collection of 140 gene-trait combinations we used to compare predictors.

A possible criticism of our predictor comparison is that UK Biobank participants carry variants that may have been used in training predictors. We considered excluding variants that had been previously reported as pathogenic or benign, e.g. in ClinVar [50], or as disease-associated or disease-causing by the Human Gene Mutation Database (HGMD) [51], as many predictors will have been trained on variants from these resources. However, we reasoned that vanishingly few variants used in predictor training sets would have been deposited in HGMD or ClinVar *on the basis of analysis of the UK Biobank data*, especially given that we have excluded common variants from our analysis. More obviously, the selection of UK Biobank participants, their genotypes, and their traits was all independent of pre-existing variant annotations. Thus, overfitting cannot be the explanation for a predictor showing improved correspondence to genotype-phenotype associations within the UK Biobank data.

Having identified the two best-performing predictors, VARITY and REVEL, we used each to guide a genome-wide burden scan. Thus, we identified 1,038 gene-trait combinations for which the burden of rare variation in the gene is correlated with case/control status for the trait, with 567 (55%) of these being novel.

Many of the novel associations seem worthy of further investigation. For example, the rare variant burden of transcription factor *ZNF226* was associated with lowering blood LDL cholesterol level, making *ZNF226* a potential target for cholesterol-lowering medications. As another example, we found the rare variant burden of *SMYD5* to be associated with lowering systolic blood pressure, suggesting *SMYD5* as a potential target for blood pressure control.

A limitation of both our predictor comparison and corresponding burden scans is that we only looked at participants of European descent in the UK Biobank cohort. Although UK Biobank is one of the largest prospective cohorts, it still contains systematic ascertainment biases (e.g. ethnicity, geography, age), so the burden test results may not be representative. Future studies could incorporate other emerging large-scale prospective cohorts to address the ascertainment biases. For example, the All of Us research program [52] supported by the National Institutes of Health (NIH) has recruited >300K US participants, of whom >50% are racial and ethnic minorities. The Grand Opportunity (GO) exome sequencing project at the National Heart, Lung, and Blood Institute (NHLBI), which contains both disease-affected and healthy participants from the US, UK and EU, could add geographic diversity. The Westlake Biobank for Chinese (WBBC) [53], which includes young Chinese participants (mean age ∼19 years), may help address the ethnicity and age-related biases in other cohorts. Also, future studies might explore mutually beneficial partnerships with private companies, e.g. clinical and direct-to-consumer genetic testing services such as Invitae, 23andMe, and commercial biobanks such as 54Gene (the first and largest for-profit genetics resource in Nigeria), to access complementary human-genetic resources.

Another limitation is that we did not adjust trait values to account for dependencies on other participant variables. For example, LDL cholesterol levels are correlated with both age and male sex [54]. In future analyses, LDL cholesterol measures adjusted for age and sex, or more precise variables such as apolipoprotein B, could have a variation that is more attributable to genetic variation and therefore show greater correlation with predictor scores. Because the same adjusted LDL cholesterol values would be used to evaluate all predictors, such an adjustment is unlikely to impact the relative rankings amongst predictors. However, using adjusted traits that correspond more closely with participant genotypes could increase the yield of burden scans.

Burden scans were performed using either a recessive or dominant model. Ideally, a recessive model would only consider a participant to have qualifying variants where both alleles carried a qualifying variant. However, as the currently available UK Biobank exome sequences are not phased, our model was not informed about whether variants were *in cis* or *in trans* with one another. Rather, the model assumed that the two most damaging variants were *in trans*. With the ongoing efforts to generate fully phased human genome sequences [55, 56], future studies might be able to improve the recessive model in burden scans by incorporating chromosomal phase information.

While this burden scan used the “qualifying variant” approach to remove the vast majority of rare missense variants that are predicted neutral, we did not account for the full spectrum of differences in variant effect. For example, PTVs close to the end of the coding region of certain genes might not contribute to a given trait as much as PTVs at the beginning of the same genes [57, 58]. A weighted burden test approach [59, 60] has been developed to assign various weights to different variant types depending on their anticipated effect on a given trait. However, this approach is limited to human traits where the effect of different variant types has been established. As our understanding of the effects of genetic variation improves (e.g. through experimentally-generated variant effect maps), it might become possible to develop a generic weighing scheme to more accurately model variant effects in the future.

Many features of this study could be directly applied to the growing body of experimentally-generated evidence about variant impacts, e.g. variant effect maps. For example, the assessment scheme we used to compare computational predictors could also be used to compare experimental variant effect maps with one another or with computational predictors. Similarly, thresholds derived from experimental variant effect maps could be used to define qualifying variants and thus to inform and potentially improve the power of burden tests.

## Conclusion

Burden scans have shown great promise for identifying disease mechanisms and therapeutic targets [39, 61]. Through an independent assessment of 20 computational variant effect predictors, this study identified two predictors, VARITY and REVEL, as corresponding most closely with human traits in a large prospective cohort, suggesting that they might also enhance the power of rare-variant burden scans for detecting gene-trait associations. Systematic burden scans performed with each of these two predictors identified many novel gene-trait combinations, including some with potential therapeutic value. Thus, variant effect predictor assessment and improved burden scan methods have offered an improved understanding of gene-trait associations for both clinical and research applications.

## Methods

### Gene-trait combinations

We assembled gene-trait combinations wherein each combination, the collection of rare missense variants in the gene has been associated with the trait by at least one of four systematic ‘burden test’ studies [28–31] of the UK Biobank cohort. From the initial set of 162 gene trait combinations, we excluded combinations for which the trait had been ascertained in fewer than 10 participants or for which the gene IDs are currently unrecognized or not linked to any proteins in Ensembl database version 104 [62]. We also excluded the *TTN* gene as non-representative given its extreme size and an enormous number of reported variants.

### Human Variants

This study was conducted with whole-exome sequencing data from 454,797 participants in the UK Biobank cohort, for which variants were retrieved from the OQFE version [63] of the whole-exome VCF files (field ID: 23148). The transfer of human data was approved and overseen by The UK Biobank Ethics Advisory Committee (Project ID [PID]: 51135). The canonical isoform of each gene product that we examined was defined according to the Ensembl database (GRCh38) [62], with exonic coding regions defined according to the CCDS database [35]. Coding variants corresponding to these coding regions were extracted from raw VCF files. Adapting the filtering criteria used by the UK Biobank [64], we examined only coding variants having a Phred quality score > 20, individual missingness < 10%, minimum read coverage depth of 7, and carried by at least one participant passed the allele balance threshold (i.e. the proportion of reads covering a variant’s location that supports the variant) > 0.15. Because evidence to support clinical interpretation is typically more abundant for common variants (e.g., from genome-wide association studies), the most critical context for computational predictors to perform well is for the rare variation. Therefore, we further restricted our analysis to rare variants, as defined here by having a global MAF < 0.1% in gnomAD [65] and the UK Biobank cohort. If a variant was not found in the gnomAD database and had MAF < 0.1% in the UK Biobank cohort, we assumed it to be rare.

### Variant effect predictions

We considered 20 computational variant effect predictors (Table S2). Scores for predictors were obtained either from a pre-existing repository [66] or by running the predictor code directly. If a predictor did not provide predictions for at least ten rare missense variants of the gene for a given gene-trait combination, it was not included in the comparison for that combination.

### Predictors comparison

We first split the 140 gene-trait combinations into two categories, depending on whether the associated traits are binary or quantitative (i.e. where measurements are continuous). Predictor comparison was conducted separately for each gene-trait combination and restricted to participants carrying rare missense variants in each gene. For example, when comparing predictions for the *LDLR*-“taking cholesterol-lowering medication” combination, we only included participants carrying rare missense *LDLR* variants.

For each variant effect predictor and gene-trait combination where the trait was binary (e.g. capturing whether or not participants were taking cholesterol-lowering medication), we rescaled predictor scores. To reduce the impact of outliers, we set the lowest 5% of scores to the 5%ile value and the highest 5% of scores to the 95%ile value. Scores were then rescaled to range from 0 to 1 (with 0 corresponding to neutral variants and 1 corresponding to damaging variants) to allow for pairwise comparison of scores from predictors with different score ranges. Variants for which the predictor failed to make predictions were not included in downstream analysis. We subsequently computed a person-centric variant score for each participant, which was calculated to be the sum of all variant scores for that participant. Continuing with the *LDLR*-“taking cholesterol-lowering medication” example above, a participant carries two missense variants in *LDLR*, each with a variant score of 0.5, will have a person-centric *LDLR* variant score of 1.

Then, participants (with their associated various person-centric variant scores) were subjected to precision vs. recall analysis. At a given score threshold (*s*), precision is defined as

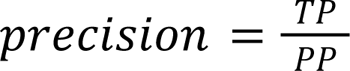

where true positives (*TP*) are participants with person-centric variant scores ≥ *s* and the trait, and predicted positives (*PP*) are participants with person-centric variant scores ≥ *s*.

Because precision depends on the prior probability (*prior*), i.e., the prevalence of the trait, we used balanced precision (the precision expected if the test set were to have an equal number of positive and negative data points) as described by Wu et al. [16]

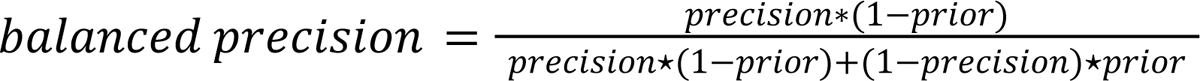

Correspondingly, recall is defined as

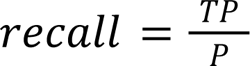

where positives (*P*) are participants with the trait.

We subsequently calculated the area under the balanced precision-recall curve (AUBPRC) using the “auprc” function in the R *yogiroc* package [67].

To better understand uncertainty in the calculated AUBPRC values, we used 1000-iteration bootstrapping (random sampling of variants with replacement) to compute a distribution of AUBPRCs for each predictor with each gene-trait combination. We then empirically determined the mean AUBPRC and the 95% confidence interval of the resampled distribution. We compared variant effect predictors in terms of mean AUBPRCs and calculated an empirical p-value for each pair of computational predictors, i.e., the fraction of pairs of resampled AUBPRC distributions, one from each predictor, in which one predictor did not achieve a higher AUBPRC than the other predictor. From the p-values, false discovery rates (FDRs) were subsequently calculated to account for multiple hypotheses testing [68]. For each gene-trait combination, predictors were ranked by performance. Any predictor that was not significantly different from the numerically best-performing predictor (using an FDR threshold of 10%) was considered tied for best.

In contrast, for gene-trait combinations with quantitative traits (e.g. blood low-density lipoprotein (LDL) cholesterol level, measured in mmol/L), we employed the Pearson correlation coefficient (PCC) to compare predictor performance. Because a variant may receive one variant effect score for each predictor but may present in multiple participants, where multiple participants carried the same variant, we averaged their quantitative trait values. This aggregation step ensured that each variant had one averaged trait value and one predicted variant effect score for a given predictor. Each variant effect predictor was examined individually. For each predictor and a given gene-trait combination, we examined variants for which the trait was measured, and a variant effect score was available.

To better understand uncertainty in the calculated PCC values, we used the same 1,000-iteration bootstrapping as documented above to compute a distribution of PCCs for each predictor with each gene-trait combination. We then empirically determined the mean PCC and the 95% confidence interval of the resampled distribution. To eliminate negative values, we used PCC^2^ instead of PCC and calculated an empirical p-value (i.e., the fraction of pairs of resampled PCC^2^ distributions, one from each predictor, in which one predictor did not achieve a higher PCC^2^ than the other predictor) for each pair of computational predictors. To account for multiple hypothesis testing, we again derived FDRs from these empirical p-values. For each gene-trait combination, predictors were ranked by performance. Any predictor that was not significantly different from the numerically best-performing predictor (FDR < 10%) was considered tied for best.

After completing the 140 predictor comparisons (one for each of the 140 gene-trait combinations), we collected the best-performing variant effect predictors from each comparison. Because multiple predictors may be tied for best (i.e. not significantly different from the numerically best-performing predictor) in a given comparison, we then ranked predictors by the number of gene-trait combinations in which the predictor was tied for best.

To assess the statistical significance of performance differences between variant effect predictors ranked above considering all gene-trait combinations, we carried out a two-tailed Wilcoxon signed-rank test comparing the number of gene-trait combinations in which a given predictor was considered best-performing (or tied for best) for each pair of predictors. FDR values for each comparison were derived as above. The same FDR threshold (FDR < 10%) was used to determine statistical significance.

### Burden testing

We conducted genome-scale burden scans, both without the benefit of computational predictors, and also separately using each of the two best-performing computational predictors, VARITY and REVEL. The burden scans cover 14,107 phenotypes (including sub-phenotypes encoded by the International Classification of Diseases, ICD-10 [69]) and rare variants (MAF < 0.1%; see above for more complete definition of rare variation) from 428,517 participants of European descent in the UK Biobank cohort. For each trait, each participant was assigned a case or control label, depending on whether participants carried a given trait (or excluded where trait status was ambiguous). For qualitative traits, this process was trivial because labels can be assigned directly. For example, participants taking cholesterol-lowering medication (a qualitative trait) were labelled as cases, and those not taking the medication were labelled as controls.

In contrast, quantitative traits needed to be converted to qualitative traits prior to label assignment. Because for some quantitative traits, such as total cholesterol level (mmol/L), measurements on both high and low extremes are considered pathological [70], we developed two conversion schemes. After ranking participants in descending order by the quantitative trait, the “top” scheme labels the top 10% participants as cases, the next 80% (10-90%) participants as controls, and excluded the remaining (bottom 10%) participants. On the contrary, the “bottom” scheme labels the bottom 10% of participants as cases, the next 80% of participants as controls and excludes the top 10% of participants. For each quantitative trait, both schemes were considered separately.

For each rare missense variant, we used VARITY and REVEL individually to compute variant effect scores. To enable efficient data processing, UK Biobank split human variants across 977 pVCF files, each containing variants within one “chromosomal region” (details on https://biobank.ctsu.ox.ac.uk/crystal/refer.cgi?id=837). Qualifying rare missense variants, i.e., variants predicted to be damaging to protein function and therefore potentially pathogenic, were therefore defined by score thresholds established separately for each trait and region using missense variants in that region. In each case, the variant score threshold used to define qualifying variants was the 95%ile of variant effect scores of variants in control participants. Rare missense variants with a variant effect score higher than the threshold were considered qualifying variants and included in burden scans. As a “no predictor” control, we also conducted a burden scan including all rare missense variants regardless of their predicted likelihood to be damaging.

To capture genetic traits with different modes of inheritance, we performed burden scans using both dominant and recessive models for the mode of inheritance. To identify participants most likely to carry damaging genetic variation in a given gene (“qualifying participants”), the *dominant* model defined qualifying participants as those carrying at least *one* qualifying variant while the *recessive* model defined qualifying participants as those carrying at least *two* qualifying variants.

For each gene-trait combination, UK Biobank participants were categorized into a 2-by-2 contingency table based on 1) whether each participant was a case or control and 2) whether each participant carried sufficient qualifying variants. Following the statistical test used in a previous state-of-the-art burden scan [39], we calculated a p-value for every gene-trait combination based on the contingency table using Fisher’s exact test [71]. To correct for multiple hypothesis testing, we computed FDR using a single-iteration permutation approach [38] used in the genome-wide burden test mentioned above [39]. The number of participants having a qualifying variant in a given gene, and the number of participants with a given trait, varied greatly between genes and traits, in some cases ranging across 3 orders of magnitude. We therefore used a null distribution for each gene-trait combination that was based on permutation-derived P-values from combinations of other genes and traits corresponding to similar numbers of participants. To that end, we divided genes into deciles based on the number of participants with a qualifying variant in that gene and divided traits into deciles based on the number of participants with that trait, thus organizing gene-trait combinations into a 10-by-10 matrix (illustrated in Figure S3). Gene-trait combinations with FDR < 5% calculated using the appropriate null distribution were considered significant.

## Supplemental data

Supplemental data includes five tables and three figures.

## Abbreviations

VUS: Variant of uncertain significance

SNV: Single-nucleotide variant

MAF: Minor allele frequency

AUBPRC: Area under the balanced precision-recall curve

FDR: False discovery rate

PCC: Pearson correlation coefficient

FID: Field ID

LDL: Low-density lipoprotein

FH: Familial hypercholesterolemia

CI: Confidence interval

BMI: Body mass index

HGMD: Human Gene Mutation Database

NIH: National Institutes of Health

GO: Grand Opportunity

NHLBI: National Heart, Lung, and Blood Institute

WBBC: Westlate Biobank for Chinese

## Declarations

### Ethics approval and consent to participate

The transfer of human data was approved and overseen by The UK Biobank Ethics Advisory Committee (Project ID: 51135).

## Consent for publication

Not applicable

## Availability of data and materials

The source code used in the variant effect predictor assessment study is available at GitHub: https://github.com/rothlab/variant-effect-predictor-assessment. The burden scan source code is available at Github: https://github.com/rothlab/burden-test-public. The UK Biobank dataset is available by application via https://www.ukbiobank.ac.uk/.

## Authors’ contributions

F.P.R., D.K. and R.L. conceived the idea and designed the study. D.K. and R.L. analyzed and interpreted the participant data from the UK Biobank cohort. Y.W. curated variant effect predictor scores for the variants examined in this study. J.W., R.A.H. and F.P.R. provided advice for the project. D.K., F.P.R. and R.A.H. wrote the manuscript. All authors read and approved the final manuscript.

## Acknowledgements

This project depended entirely on UK Biobank participants, and on those who envisioned, supported, developed and maintain this amazing resource. We are grateful to Jennifer Knapp and Dayag Sheykhkarimli for their helpful comments and Thomas Hu and Jeff Liu for computational support.

