## Supplementary material for "Empowering rare variant burden-based gene-trait association studies via optimized computational predictor choice": Figure S1-3

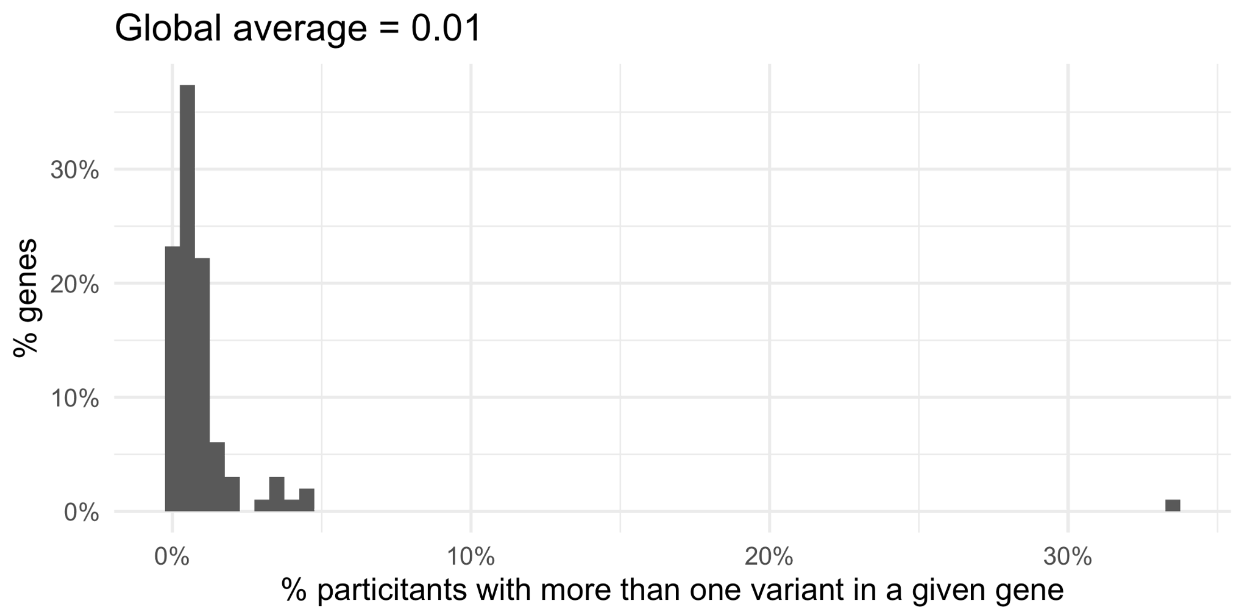


**Figure S1.** The fraction of participants with more than one variant in their gene for each of the 99 genes in the 140 gene-trait combinations.

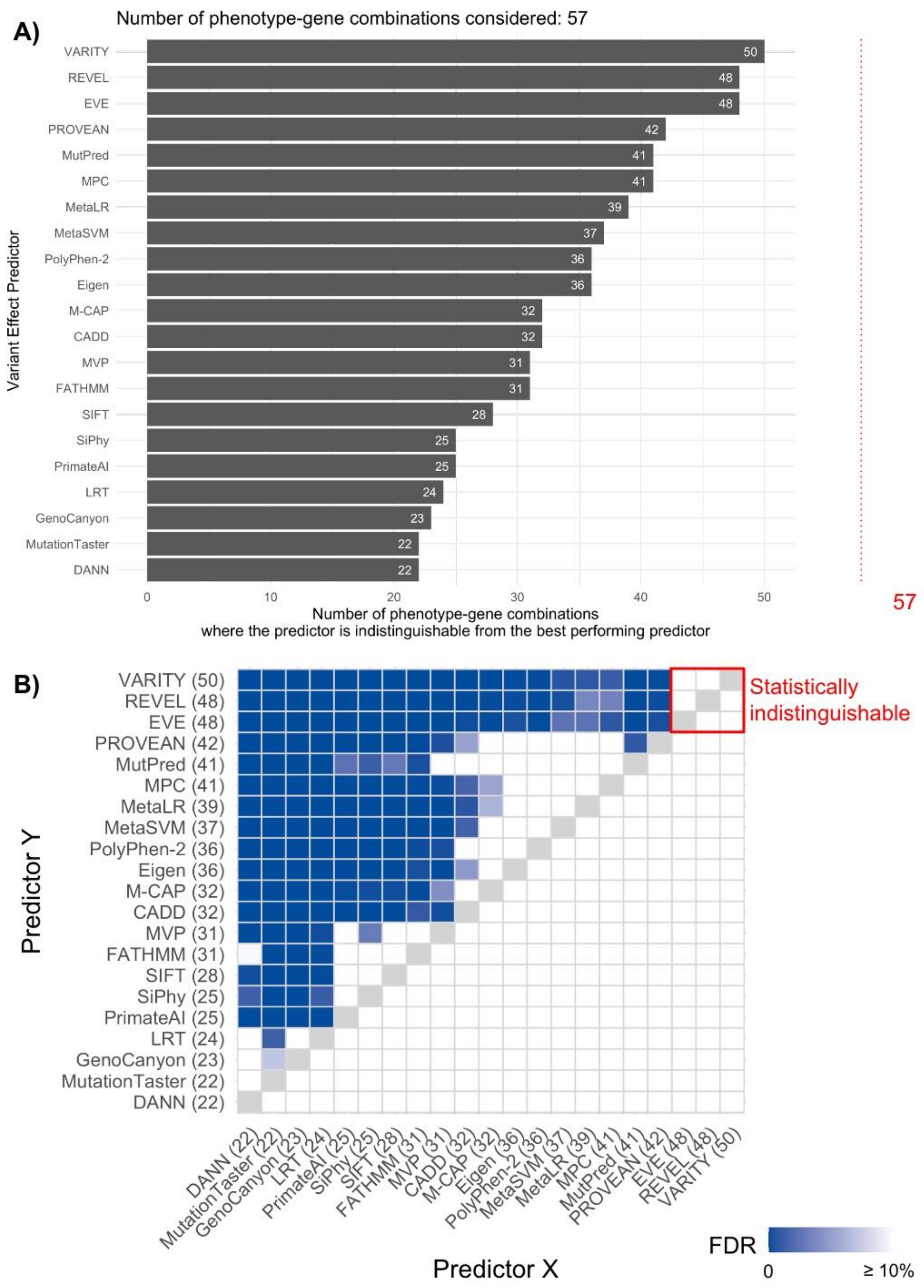


**Figure S2. A)** The number of gene-trait combinations for which each predictor is either best-performing or indistinguishable from best-performing. The red line highlights the maximum value based on the 57 combinations where all 20 predictors could make enough predictions. **B)** Comparisons between all pairs of computational predictors, determining if the performance of one predictor is significantly different than another. Variant effect predictors are ranked top-to-bottom and right-to-left based on decreasing number of gene-trait combinations where the predictor is among the best-performing predictors. Comparisons in which one predictor Y is significantly different from another predictor X (with FDR < 10%) are indicated in blue.

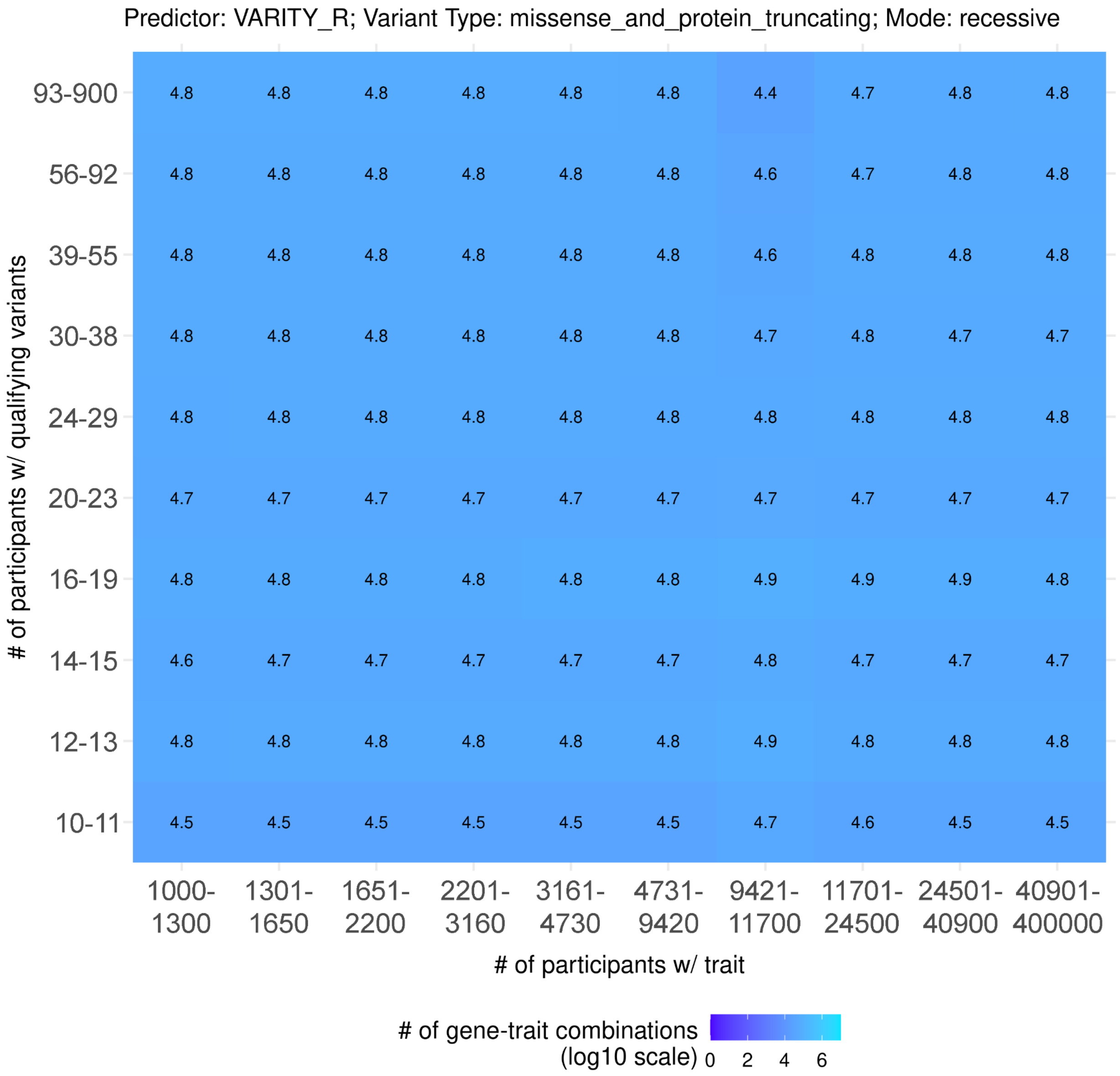


**Figure S3.** The number of gene-trait combinations in a 10-by-10 grid. Genes were divided into deciles based on the number of participants with a qualifying variant in that gene and traits were divided into deciles based on the number of participants with that trait.
